# Macrophages orchestrate the expansion of a pro-angiogenic perivascular niche during cancer progression

**DOI:** 10.1101/2020.10.30.361907

**Authors:** James W. Opzoomer, Joanne E. Anstee, Isaac Dean, Emily J. Hill, Ihssane Bouybayoune, Jonathan Caron, Tamara Muliaditan, Peter Gordon, Dominika Sosnowska, Rosamond Nuamah, Sarah E. Pinder, Tony Ng, Francesco Dazzi, Shahram Kordasti, David R. Withers, Toby Lawrence, James N. Arnold

## Abstract

Tumor associated macrophages (TAMs) are a highly plastic stromal cell type which support cancer progression. Using single-cell RNA-sequencing of TAMs from a spontaneous murine model of mammary adenocarcinoma (*MMTV-PyMT*) we characterize a subset of these cells expressing lymphatic vessel endothelial hyaluronic acid receptor 1 (Lyve-1) which spatially reside proximal to blood vasculature. We demonstrate that Lyve-1^+^ TAMs support tumor growth and identify a pivotal role for these cells in maintaining a population of perivascular mesenchymal cells which express alpha-smooth muscle actin and phenotypically resemble pericytes. Using photolabeling techniques show that mesenchymal cells maintain their prevalence in the growing tumor through proliferation and uncover a role for Lyve-1^+^ TAMs in orchestrating a selective platelet-derived growth factor-CC-dependent expansion of the perivascular mesenchymal population, creating a pro-angiogenic niche. This study highlights the inter-reliance of the immune and non-immune stromal network which support cancer progression and provides therapeutic opportunities for tackling the disease.

## Main text

Tumor associated macrophages (TAMs) form a major part of the stromal cell infiltrate in solid tumors (*1*), and are highly plastic to their environment which creates phenotypic and functional diversity within the population (*2, 3*). Tumors exploit the plastic nature of TAMs to facilitate disease progression through promoting angiogenesis (*4, 5*), immune suppression (*6, 7*), chemotherapeutic resistance (*8–10*) and tumor cell migration and metastasis (*2, 11–15*). Although macrophage polarization has a spectrum of possible phenotypes that can be adopted (*16, 17*), it is apparent that functionally important subsets preferentially accumulate, that are guided by spatial and environmental cues, to conduct specialized tasks vital to tumor progression (*2, 6, 17–20*). To resolve the heterogeneity of the TAM population, CD45^+^Ly6G^-^CD11b^+^F4/80^hi^ TAMs were FACs cell-sorted from enzyme-dispersed tumors from *MMTV-PyMT* mice (*21*) (fig. S1A). The TAMs were then subjected to the droplet-based 10X Genomics Platform for single cell RNA-sequencing (scRNA-seq; Fig. 1A). A total of 9,039 TAMs were sequenced across three individual tumors. Unsupervised graph-based clustering of the transcriptomes, visualized using *UMAP* (*22*), revealed eight distinct transcriptomic TAM clusters (Fig. 1B-D and fig. S1B,C). The presence of these transcriptomic clusters, despite the tumors being spontaneous, were conserved across the three tumors analyzed (Fig. 1E). Gene Ontology (GO) analysis of the transcriptional programs within these clusters revealed diversity in both the number and type of biological pathways that were active. One cluster (TAM08) represented a highly proliferative TAM state, indicating that TAMs are capable of proliferation in the tumor microenvironment, however these TAM’s transcriptome was dominated singularly by cell-cycle associated genes and so was not carried forward for further functional analysis (Fig. 1F and fig. S1D). Interestingly, the TAM clusters with few enriched GO terms, that appeared to be the least polarized in their gene expression profile (TAM01 and 02), represented almost a quarter of TAMs within the tumor (23.3% ± 3.4 of all TAMs analyzed), suggesting that a significant proportion of TAMs remain relatively unspecialized in their role (Fig. 1E,F and fig. S1E). Trajectory analysis using *Slingshot* (*23*) and diffusion maps was able to align the 8 identified clusters into a three trajectory polarization model with TAM04, 06 and 07 clusters representing predicted polarization extremes (Fig. 1G,H and fig. S2). Analysis of the three developmental pathways for their enrichment of M1/M2 (*24*) programs using the marker gene list of Orecchioni *et al* (*25*) highlighted TAM04 to be skewed towards an inflammatory (M1-like) transcriptome (Fig. 2A,B) which were more enriched for expression of inflammatory genes representative of a cellular response to type-1 interferons such as *Irf7* and *Isg15*. TAM06 and TAM07 possessed a more pro-tumoral (M2-like) transcriptome (Fig. 2A,B). TAM06 was more enriched for anti-inflammatory genes such as *Il10*, whereas both TAM06 and TAM07 were enriched in *Ccl2*, *Mmp19*, *Hb-egf* and also *Mrc1* (the gene for MRC1/CD206) (*26*). However, TAM06 and TAM07 were functionally distinct in many of their enriched GO biological pathways, with a preferential skewing of TAM06 towards angiogenic processes and TAM07 towards immune regulation, highlighting a specialized sub-division of roles within the tumor (Fig. 2C,D). Flow cytometry analysis of gated CD206^+^F4/80^hi^ TAMs stained for markers identified within the scRNA-seq analysis, confirmed that similar TAM sub-populations could be distinguished using the predicted protein markers in *MMTV-PyMT* tumors. The CD206 expressing pro-tumoral TAMs could be differentiated based on their expression level of CD206, MHCII, and the lymphatic vessel endothelial hyaluronic acid receptor 1 (Lyve-1) (Fig. 2E,F), into CD206^lo^MHCII^lo^Lyve-1^-^ (TAM05) and the predicted pro-tumoral polarization extremes of CD206^hi^MHCII^lo^Lyve-1^+^ (Lyve-1^+^ TAMs; TAM06) and CD206^int^MHCII^hi^Lyve-1^-^ (TAM07). Lyve-1 has traditionally been considered a marker of lymphatic endothelium (*27*), but has also been utilized as a marker on tissue-resident macrophages (*28–32*) and TAMs (*33*). The Lyve-1^+^ TAM subset (TAM06) accounted for 10.7±3.5% of total TAMs and 1.4±0.4% of live cells within the tumor (Fig. 2G).

**Figure 1.**
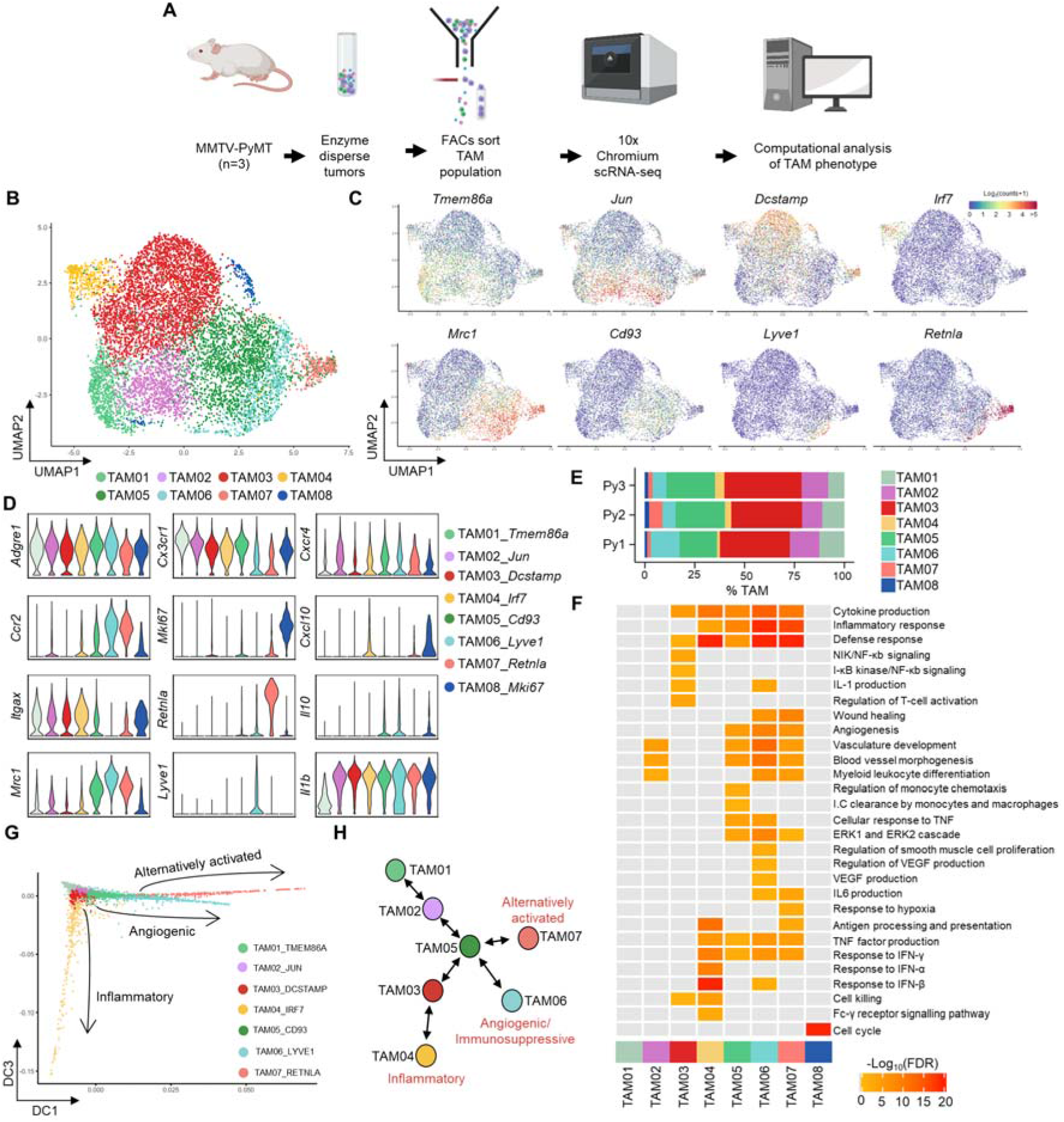
ScRNA-seq of TAMs in *MMTV-PyMT* tumors reveals three distinct polarization pathways. (**A**) Schematic outlining the scRNA-seq experimental workflow which was conducted for n=3 individual *MMTV-PYMT* tumors and mice, sequencing a total of 9,039 cells using the 10X Genomics’ Chromium platform. (**B**) UMAP plot of sequenced TAMs colored by their associated cluster identity. (**C**) UMAP visualizations of predicted marker gene expression for distinct TAM clusters shown in (**B**). (**D**) Violin plots of selected genes associated with TAM cluster identity seen in (**B**). (**E**) The relative proportion of each TAM cluster across the individual *MMTV-PyMT* tumors analyzed. (**F**) Heatmap representing significantly upregulated GO pathway terms in one or more TAM clusters. (**G,H**) Scatter plot of single cells projected into two dimensions using diffusion maps, where each cell (dot) is colored by cluster identity, labeled with diffusion component (DC) space annotation representing lineage trajectories predicted by the *Slingshot* package (**G**) and schematic map of each TAM cluster’s location along the respective trajectories (**H**).

**Figure 2.**
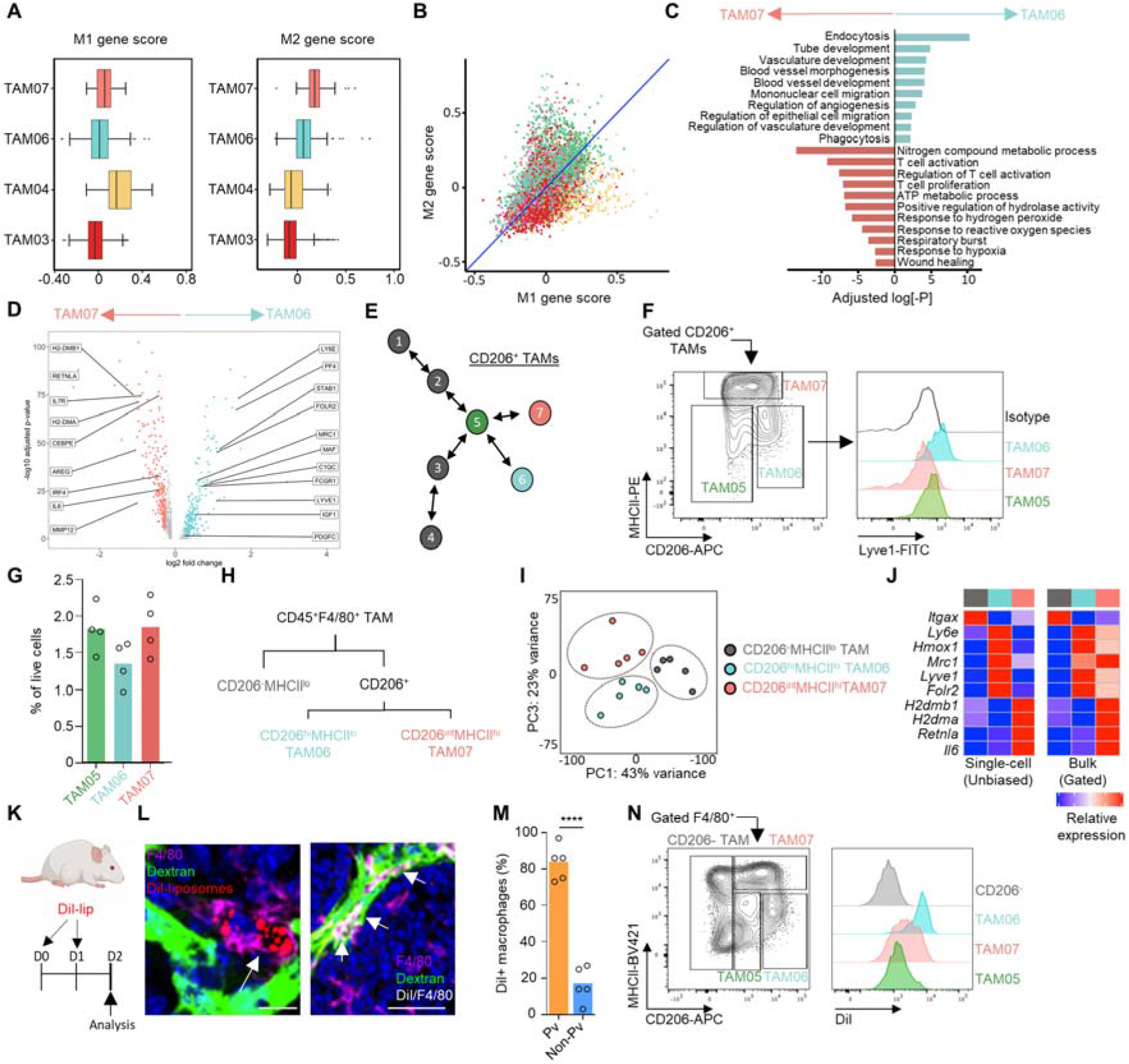
Lyve-1 marks a pro-tumoral perivascular TAM population. (**A,B**) TAM clusters identified in Fig.1 using scRNA-seq (n=3 mice) were assessed for their similarity to M1/2 macrophage polarization programs. (**A**) Box and whisker plots show normalized mean M1 and M2 associated gene scores across the indicated TAM clusters identified. (**B**) Scatter plot of normalized mean M1/M2 gene score plotted by individual cell (dot) and colored according to their respective TAM cluster. Blue line represents y=x line for reference. (**C,D**) Subset unique, significantly upregulated GO terms based on differentially expressed genes between the two terminal *Mrc1*^high^ TAM subsets identified in the TAM scRNA-set dataset (**C**), volcano plot showing differentially expressed genes between the two subsets of pro-tumoral TAM (**D**). (**E-G**) Schematic of *Slingshot* trajectory analysis of TAM clusters highlighting predicted *Mrc1* expressing clusters. The clusters where *Mrc1* was not identified as a differentially expressed gene are greyed out (**E**), and mapping of these clusters predicted by the scRNA-seq dataset onto a contour plot of FACs-gated live (7AAD^-^) CD206^+^ F4/80^hi^ TAMs from enzyme-dispersed *MMTV-PyMT* tumors. TAM populations are separated based on their respective expression of CD206 and MHCII (left panel) and then assessed for their expression of Lyve-1, shown as histograms (right panel; colored shaded histograms) against that of the isotype control staining of F4/80^+^ TAMs (open black line) (**F**) and quantification of the gated populations (**G**). Data representative of n=4 tumors. (**H-J**) Diagram of the FACs gating strategy for TAM populations sorted for bulk RNA-seq (n=5 tumors) (**H**), PCA plot of the bulk-sequenced TAM populations using the top 2,000 most variable genes (**I**) and heatmaps comparing the relative expression of selected differentially expressed genes for TAM clusters (left), population color is indicative of the populations identified in (**H**,**I**) and isolated TAM populations subjected to bulk RNA-seq (right panel), (**J**). (**K-N**) Schematic for experimental approach and dosing strategy to label PvTAMs using Dil-labeled liposomes (**K**). Representative images of frozen sections of *MMTV-PyMT* tumors showing DAPI (nuclei; blue), i.v. dextran marking vasculature (green), Dil from the liposomes (red) and antibody staining against F4/80 (magenta), left shows an example Dil-labeled TAM and right panel shows a larger tumor area displaying co-localizing pixels for Dil and F4/80 as white. White arrows highlight example pvTAM cells which have been labeled by Dil-containing liposomes (**L**) Scale bar 25μm (left panel) and 50μm (right panel). Quantification of the spatial location of Dil^+^ F4/80^+^ TAMs relative to vasculature (Pv-perivascular) across multiple sections in each tumor across n=5 mice (**M**). Analysis of the TAM population phenotype up-taking Dil-liposomes from enzyme-dispersed tumors within the F4/80^hi^ CD206^+^ subsets – gate as shown (left) and histogram of the indicated TAM subsets Dil fluorescence (right) (**N**). Box and whisker plots, the boxes show median and upper and lower quartiles and whiskers shows the largest value no more than 1.5*IQR of the respective upper and lower hinges, outliers beyond the end of the whisker are plotted as individual dots. Bar charts represent mean and the dots show individual data points from individual tumors and mice. ** *P*<0.01, *** *P*<0.001.

To validate that the populations identified in the scRNA-seq and flow cytometry data were equivalent, the FACs-gated populations were subjected to bulk population RNA-seq alongside CD206^-^MHC^lo^F4/80^hi^ TAMs as a comparator group. Principal component (PC) analysis confirmed these populations to be transcriptionally distinct (Fig. 2H,I). Comparing the bulk population RNA-seq to that of the scRNA-seq populations validated close concordance between the identified populations across a range of predicted marker genes (Fig. 2J). Lyve-1^+^ TAMs (TAM06) also selectively expressed the transcription factor *Maf* (fig. S2D) and CD206^int^MHCII^hi^Lyve-1^-^ (TAM07) the transcription factor *Retnla* (Fig. 1D), which may indicate that these transcription factors play a role in polarization identity. A monocyte-derived macrophage with a similar MHCII^lo^Lyve-1^hi^ surface phenotype has been demonstrated to reside proximal to vasculature in a variety of healthy tissues (*32*). GO pathway analysis also suggested that Lyve-1^+^ TAMs were highly endocytic (Fig. 2C). Liposomes containing the fluorescent lipophilic dye 1’-dioctadecyl-3,3,3’,3”tetramethylindocarbocyanine perchlorate (Dil) have previously been used to study perivascular TAM (pvTAM) development (*13*) and we predicted they could represent a tool to preferentially label the Lyve-1^+^ TAM subset. We developed a labeling protocol that could selectively mark pvTAMs (fig. S3A). Confocal analysis of the tumors demonstrated that Dil-liposomes labeled a population of pvTAMs (Fig. 2K-M) and *ex vivo* characterization of the Dil-labeled cells in enzyme-dispersed tumors confirmed their phenotype to indeed be that of the Lyve-1^+^ TAM subset (TAM06, Fig. 2N).

As the liposome labeling protocol preferentially labeled Lyve-1^+^ TAMs (Fig. 2M,N and fig. S3A-B), we utilized clodronate-filled liposomes (*34*) under an equivalent administration protocol as a means to selectively deplete the population and investigate their possible role in tumor progression. Depletion of these cells in *MMTV-PyMT* tumors resulted in a significant slowing of tumor growth (Fig. 3B), highlighting a fundamental role for these cells in tumor progression. Even over the long-term administration of clodronate-filled liposomes, which displayed little sign of toxicity in the animals (fig. S3C), provided a preferential depletion of Lyve-1^+^ TAMs (TAM06), largely sparing the CD206^int^MHCII^hi^Lyve-1^-^ (TAM07) subset of specialized pro-tumoral TAMs (Fig. 3C-E), CD206^-^ TAMs (Fig. 3F), and CD11b^+^Ly6C^+^ monocytes (Fig. 3G). Furthermore, using immunofluorescence imaging there was an observable selective spatial loss of perivascular TAMs (pvTAMs) within the clodronate-filled liposome treated mice (Fig. 3H), where the majority of TAMs surrounding blood vessels were no longer observable. To understand the mechanism through which Lyve-1^+^ pvTAMs promote tumor progression (Fig. 3B), we first phenotyped the immune-infiltrate of the tumors. Loss of Lyve-1^+^ pvTAMs did not change the abundance of any immune cell populations analyzed within the tumor microenvironment, other than a statistically significant increase in the abundance of the migratory CD11c^+^CD103^+^dendritic cells (DCs) (Fig. 3I and fig. S3D), which contribute to cytotoxic T-lymphocyte recruitment in the tumor (*35*) and priming of the anti-tumor immune response (*36*). However, there was no increase in CD8^+^ or CD4^+^ T-cell recruitment post depletion of Lyve-1^+^ pvTAMs (Fig. 3I). Perivascular macrophages are known to play a role in angiogenesis (*18*), and the Lyve-1^+^ TAM population expressed pathways associated with angiogenesis (Fig. 2C), which could account for the control of tumor growth observed when the TAM subset was depleted (Fig.3B). Immunofluorescence analysis of these tumors had shown no overall change in density of endothelial cells within the tumor (Fig. 3K), but the tumors themselves were smaller (Fig.3B). Further analysis of sections from *MMTV-PyMT* tumors stained for CD31^+^ endothelial cells and perivascular α-smooth muscle actin (αSMA) expressing stromal cells revealed a change in vessel architecture (Fig. 3J), where depletion of Lyve-1^+^ TAMs resulted in an increase in the number of individual vessel elements in the tumor (fig. S3E) with the vessel elements appearing smaller and less branched (Fig.3J). However, most strikingly, there was a loss of αSMA^+^ stromal cells proximal to vasculature (Fig. 3J,L), highlighting a potential role of Lyve-1^+^ TAMs in maintaining this stromal population.

**Figure 3.**
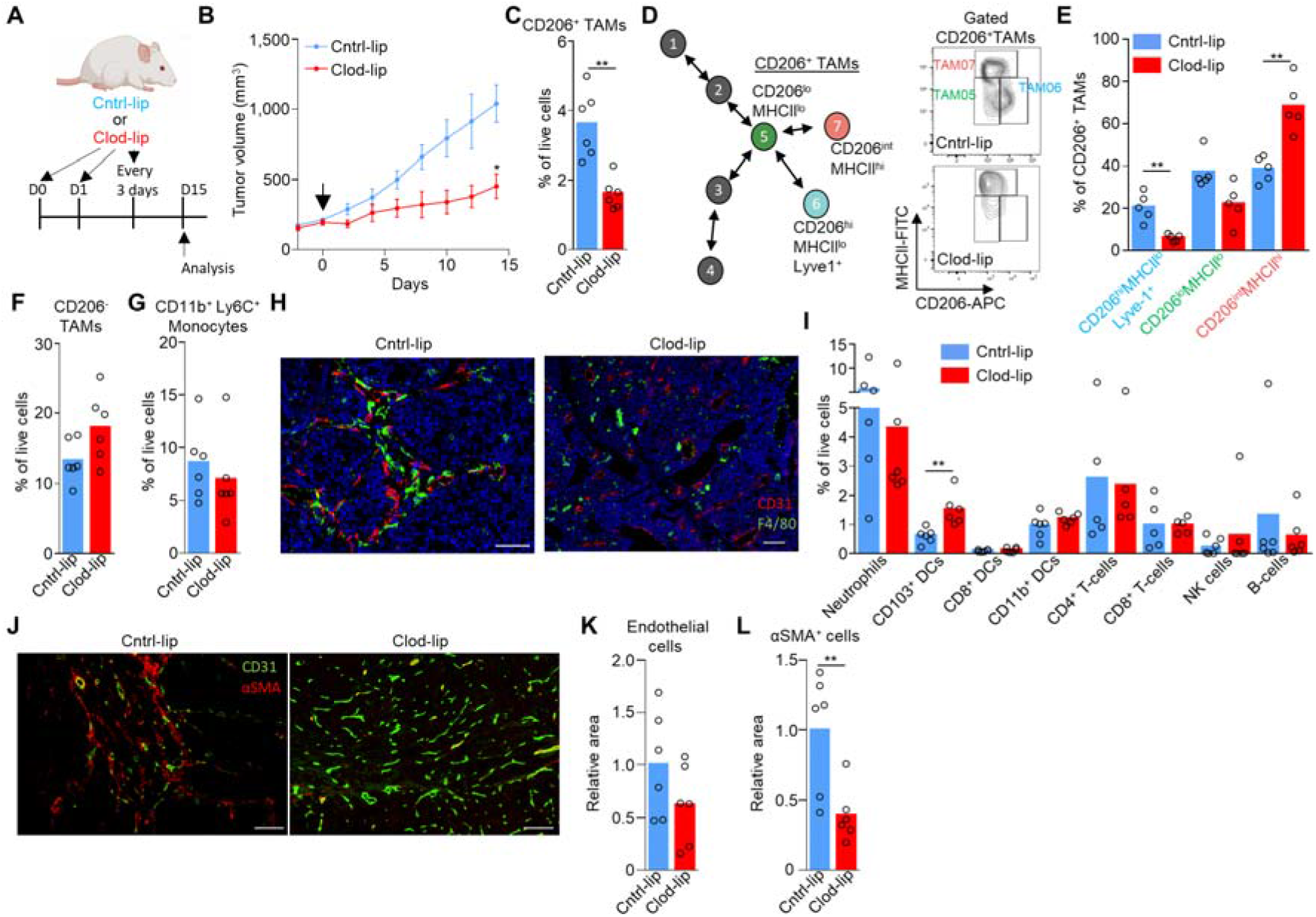
Lyve-1^+^ pvTAM depletion slows tumor growth and is associated with a concurrent loss of perivascular αSMA^+^ stromal cells. (**A**) Schematic for experimental approach and dosing strategy to deplete Lyve-1^+^ TAMs using clodronate-filled liposomes. Arrows represent days of treatment. (**B**) Growth curves of *MMTV-PyMT* tumors in mice treated with control PBS-filled liposomes (Cntrl-lip) or clodronate-filled liposomes (Clod-lip) as shown in panel (**A**), arrow marks the initiation of treatment, (cohorts of n=6 mice). (**C-I**) Tumors were excised at day 15 post initiation of administration of clodronate-filled liposomes shown in (**B**), enzyme-dispersed and assessed using flow cytometry (n=5-6 tumors in each condition) for; (**C**) Abundance of live (7AAD^-^) CD45^+^Ly6C^-^F4/80^+^CD206^+^ TAMs from enzyme-dispersed *MMTV-PyMT* tumors measured by flow cytometry. (**D**) Schematic of CD206^+^ TAM clusters identified in scRNA-seq (left in color), and a representative flow cytometry contour plots showing both cntrl- and clodronate-filled liposome treated tumors demonstrating depletion of CD206^hi^MHCII^lo^(Lyve-1^+^) TAMs within the CD45^+^Ly6C^-^F4/80^+^ TAM gate (right) and (**E**) their quantification. (**F**) Abundance of live (7AAD^-^) CD45^+^Ly6C^-^F4/80^+^CD206^-^ TAMs and (**G**) live CD45^+^CD11b^+^Ly6C^+^ monocytes. (**H**) Representative images of frozen sections of tumor sections from mice treated with cntrl- or clodronate-filled liposomes stained with DAPI (nuclei; blue) and antibodies against F4/80 (green) and CD31 (red). Scale bar represents 50μm (left panel) and 100μm (right panel). (**I**) The abundance of major immune cell types in the tumor microenvironment measured by flow cytometry. (**J-L**) Representative images of frozen sections of *MMTV-PyMT* tumors from mice treated with cntrl- or clodronate-filled liposomes stained with antibodies against CD31 (green) and αSMA (red), scale bar represents 100μm (left and right panels) (**J**) and the quantification of relative CD31^+^ pixel area (**K**) and αSMA^+^ pixel area (**L**) A total of n=12 sections were analyzed across the 6 tumors in each cohort. Growth curve in (**B**) is presented as mean ± s.e.m and bar charts represent mean and the dots show individual data points from individual tumors and mice. * *P*<0.05, ** *P*<0.01, \*\*\**P*<0.001.

Staining tissue sections from *MMTV-PyMT* tumors for the αSMA^+^ cells and F4/80^+^ TAMs placed these populations in a close spatial arrangement providing opportunity for interactions and suggested a ‘niche’ formation (Fig. 4A). This co-localization was also evident in human invasive breast carcinomas, where CD68^+^ TAMs and αSMA^+^ cells could be found in close proximity adjacent to CD31^+^ endothelial cells lining blood vessels (Fig. 4B). Interestingly, this relationship was not observed in ductal carcinoma *in situ* (DCIS), where the only αSMA^+^ cells, likely to be myoepithelial cells, appeared surrounding ductal structures and did not associate frequently with CD68^+^ TAMs. This suggests that the spatial arrangement could be associated with progressive disease where there is ongoing neoangiogenesis (Fig. 4B). To further investigate these perivascular Lyve-1^+^ TAM-dependent αSMA^+^ cells, we characterized the heterogeneity of a broad pool of tumor-associated mesenchymal stromal cells (collectively termed cancer associated fibroblasts; CAFs) using flow cytometry within enzyme-dispersed *MMTV-PyMT* tumors. The CD45^-^CD31^-^CD90^+^ population accounted for 4.0±1.6% of total live cells within 350mm^3^ tumors and their abundance increased as tumors progressed (Fig. 4C). We screened the CD45^-^CD90^+^ population for cell surface markers associated with mesenchymal subsets, including; Ly6a, CD34, PDGFRα, FAP and CD29 (*37–41*). Clustering of the multi-parametric flow cytometry data using *UMAP* (*22*) and FlowSOM (*42*) distinguished two distinct subsets (Fig. 4D). The first subset ‘CAF1’ was CD29^hi^CD34^-^Ly6a^-^FAP^lo^PDGFRα^lo^ and the second ‘CAF2’ was CD29^lo^CD34^+^Ly6a^+^FAP^hi^PDGFRα^hi^ (Fig. 4D). The two populations were FACs-sorted based on their differing expression of CD34 (Fig. 4E–F) for bulk RNA-seq to identify the αSMA-expressing population. This analysis demonstrated clear transcriptional differences in these subsets (Fig. S4A-C). The CD34^+^ CAF population displaying an inflammation-related program (fig. S4C) while the CD34^-^ CAF population expressed high levels of αSMA (*Acta2*) (Fig. 4F) and displayed a broader extracellular matrix/angiogenesis-related program (fig.S4C). These were largely similar to the CAF subsets identified in pancreatic ductal adenocarcinoma (PDAC) (*37*), however there were also key differences such as *Il6* was not a discriminatory marker for the CAF populations in *MMTV-PyMT* tumors (Fig. 4F). Interestingly, the CD34^-^ CAF population also expressed *Des, Pdgfrb* and *Cspg4* (Fig. 4F) which are genes that are often associated with pericytes, a population of specialized vessel-associated cells (*43, 44*). To confirm the presence of pericyte markers desmin (*Des*), PDGFRβ (*Pdgfrb*) and NG2 (*Cspg4*) at the protein level in these cells, immunofluorescence staining of tissues sections from *MMTV-PyMT* mice confirmed that the perivascular αSMA^+^ cells also were desmin^+^ (fig. S4D), and *ex vivo* flow cytometry confirmed the presence of surface PDGFRβ and NG2 (fig. S4E,F). CD34^-^ CAFs expressed PDGFRα, albeit low relative to the CD34^+^ population (Fig. 4D), which is regarded as a broad marker of fibroblasts. However, the pericyte marker NG2 and fibroblast marker PDGFRα colocalized at the protein level on these cells (fig. S4F), suggesting the population may represent either a ‘pathological’ pericyte phenotype or a pericyte-like CAF population. The CD34^-^ CAF population also displays similarities in gene expression to vasculature-associated ‘vCAFs’ recently characterized in MMTV-PyMT tumors, although vCAFs did not have detectable surface protein expression of NG2 (*40*). A phenotypically similar pericyte-like CAF population expressing CD29, PDGFRβ and high levels of αSMA, has also been identified in human breast cancer (*38*). Heterogeneous expression of CD34 differentiated CAF populations across different ectopic tumor models including B16, LL2 and orthotopic 4T1 (fig. S4G,H). Due to αSMA representing a defining feature of these cells we elected to refer to these cells herein as ‘αSMA^+^ CAFs’. Analyzing the abundance of the CAFs populations over the different stages of tumor progression from the healthy mammary gland, hyperplasia and the growing tumor revealed a relative increase in the abundance of the αSMA^+^ CAFs within the broader CAF population over tumor progression, suggesting a preferential selection of this subset within the tumor microenvironment (Fig. 4G). To elucidate the route through which these cells were accumulating in the tumor, we first explored local proliferation and pulsed mice bearing *MMTV-PyMT* tumors with 5-ethynyl-2’-deoxyuridine (Edu) to label actively proliferating cells. Although both CD34^+^ CAF and αSMA^+^ CAF populations displayed evidence of proliferation by comparison with healthy mammary gland, the αSMA^+^ CAFs were proliferating at a significantly faster rate (Fig. 4H). To address whether the proliferation was sufficient to account for their preferential expansion with tumor growth we utilized the *Kaede* mouse (*45*) crossed to the *MMTV-PyMT* model. Using this approach, we were able to photoconvert all tumor and stromal cells within a 100mm^3^ tumor from *Kaede*-green to *Kaede*-red (Fig. 4I). Analyzing tumors 72h after photoconversion demonstrated that CD45^+^ stromal cells predominantly displayed *Kaede*-green, highlighting the continual recruitment of hematopoietic stromal cells to the tumor from the periphery (*13, 46, 47*). In contrast, both CD34^+^ CAFs and αSMA^+^ CAF populations remained *Kaede*-red, which indicated that both CAF populations derived from a tumor-resident source of cells and was not dependent on recruitment (Fig. 4I). Therefore, the rapid proliferation of the αSMA^+^ CAFs relative to CD34^+^ CAFs may also contribute to the dynamics of CAF heterogeneity over tumor growth (Fig. 4G).

**Figure 4.**
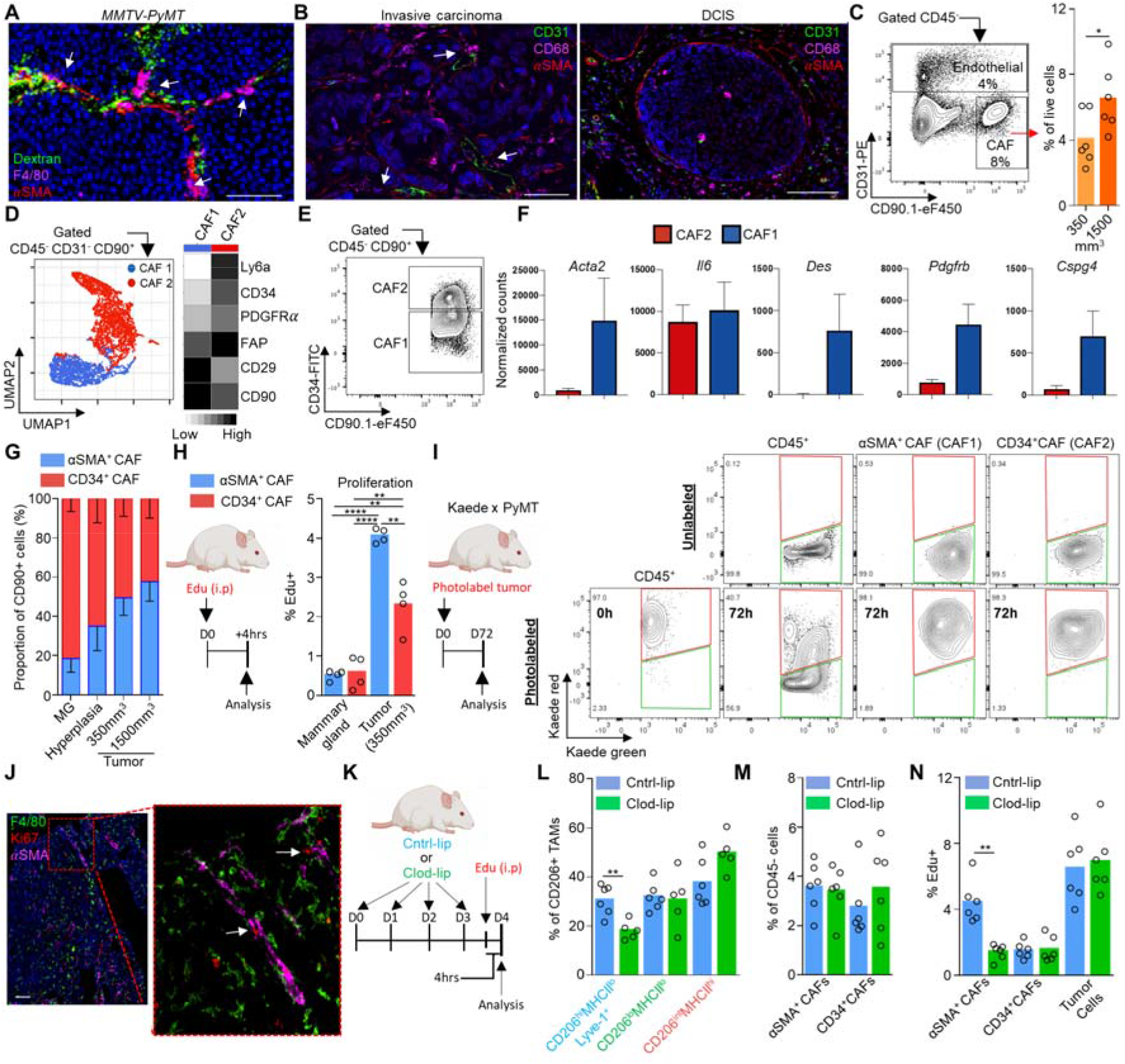
Lyve-1^+^ TAMs orchestrate pericyte-like αSMA^+^ CAF expansion within the perivascular niche of the tumor. (**A**) Representative images of frozen sections of *MMTV-PyMT* tumors stained with DAPI (nuclei; blue) and antibodies against F4/80 (magenta) and αSMA (red); functional vasculature was labeled *in vivo* using i.v. dextran-FITC (green). Scale bar represents 100μm. (**B**) Representative images of frozen sections from human invasive ductal mammary carcinoma (left) and DCIS (right) stained with DAPI (nuclei; blue) and antibodies against CD31 (green), CD68 (magenta) and αSMA (red), images representative of 5 patients. Scale bar represents 100μm (left panel) and 50μm (right panel). (**C**) Representative flow cytometry gating strategy for live (7AAD^-^) CD45^-^ cells and CD31^+^ endothelial cells and CD90^+^ CAFs (left) and the abundance of CAFs at different tumor volumes (right), n=6 mice per condition. (**D**) Identification of CAF subsets by unsupervised clustering from multiparametric flow cytometry data using the *FlowSOM* algorithm. UMAP and unsupervised clustering was performed using the markers shown in the heatmap (right). UMAP plot shows individual cells colored by their unsupervised clustering assignment (left). Heatmap displays the relative marker expression of each marker among the two subsets (right), n=4 mice. (**E**) Representative gating strategy for flow cytometry sorting the predicted subsets of CAFs by unsupervised clustering analysis. (**F**) Bar plots depicting normalized gene expression values for the indicated genes in the two bulk RNA-sequenced CAF subsets (across n=5 mice), showing that the αSMA^+^ CAF population expresses pericyte markers (*Acta2, Des, Pdgfb and Cspg4*) in *MMTV-PyMT* tumors. (**G**) Abundance of the respective CAF populations during distinct stages of tumor progression, n=6 mice per stage. (MG; mammary gland). (**H**) Schematic for experimental approach and dosing Edu into *MMTV-PyMT* mice to assess *in vivo* proliferation (left), proportion EdU^+^ cells within each CAF subset (right). (**I**) Established tumors in *Kaede MMTV-PyMT* mice were photoconverted to kaede red and then at 72h post photoconversion tumors were analyzed (schematic left) for their respective kaede red/green proportion using flow cytometry for evidence of peripheral recruitment (Kaede green cells). A representative unconverted tumor is shown for comparison (right top). (**J**) Representative images of frozen sections of *MMTV-PyMT* tumors stained with antibodies against F4/80 (green), αSMA (magenta) and the proliferation marker Ki67 (red). White arrows show αSMA^+^Ki67^+^ cells in contact with F4/80^+^ TAMs. (**K**) Schematic for experimental approach and dosing strategy to acutely deplete Lyve-1^+^ pvTAM with clodronate-filled liposomes. (**L**) Abundance of TAM CD206^+^TAM populations following cntrl- or clodronate-filled liposome treatment (n=6 mice cntrl-lip and n=5 mice clod-lip). (**M**) Abundance of CD45^-^ cell populations (cohorts of n=6 mice). (**N**) Proportion of EdU^+^ cells within each CD45^-^ cell subset, (cohorts of n=6 mice). Bar charts represent mean, error bars represent s.d. and the dots show individual data points from individual tumors and mice. * *P*<0.05, ** *P*<0.01, \*\*\**P*<0.001.

Immunofluorescence analysis for Ki67, a marker of proliferation (*48*), on αSMA^+^ cells, which we had identified as perivascular, confirmed a close spatial relationship between proliferating Ki67^+^αSMA^+^ CAFs and F4/80^+^ TAMs (Fig. 4J). To investigate whether Lyve-1^+^ pvTAMs might be implicated in the expansion of αSMA^+^ CAFs, we analyzed the incorporation of Edu after the depletion of Lyve-1^+^ pvTAMs using clodronate-filled liposomes (Fig. 4K,L). Despite no observable drop in the proportion of αSMA^+^ CAFs within the tumor over the short-term acute treatment regimen (Fig. 4M), in the absence of Lyve-1^+^ pvTAMs, high rate of proliferation of the αSMA^+^ CAF population was significantly diminished, while proliferation of the CD34^+^ CAF and tumor cell compartments remained unaffected (Fig. 4N).

To resolve how Lyve-1^+^ pvTAMs could be orchestrating αSMA^+^ CAF expansion within the perivascular niche, we utilized *CellPhoneDB*, a manually curated repository and computational framework to map the possible biological ligand:receptor interactions within RNA-seq datasets (*49*) between the Lyve-1^+^ pvTAMs, αSMA^+^ CAFs and CD31^+^ endothelial cells (which were all bulk-population RNA-sequenced) to construct an interactome of the major cell types in the perivascular niche (Fig. 5A). There were a total of 653 possible unique ligand:receptor interactions between these three cell types, highlighting the range of potential crosstalk between these populations in constructing the perivascular niche (fig. S5A). To refine this list, we selected for known mitogenic non-integrin mediated ligands which were enriched in Lyve-1^+^ pvTAMs compared to other TAM populations and could interact with receptors specifically expressed on αSMA^+^ CAFs and not endothelial cells (Fig. 5B,C). This highlighted the selective crosstalk between these two proximal cells involving *Pdgfc* (*50*) expressed by the Lyve-1^+^ pvTAMs signaling to *Pdgfra* on the αSMA^+^ CAFs within the perivascular niche (Fig. 5C). More broadly, the Lyve-1^+^ TAM subset was a major source of *Pdgfc* in the tumor (Fig. 5D and S5B).

**Figure 5.**
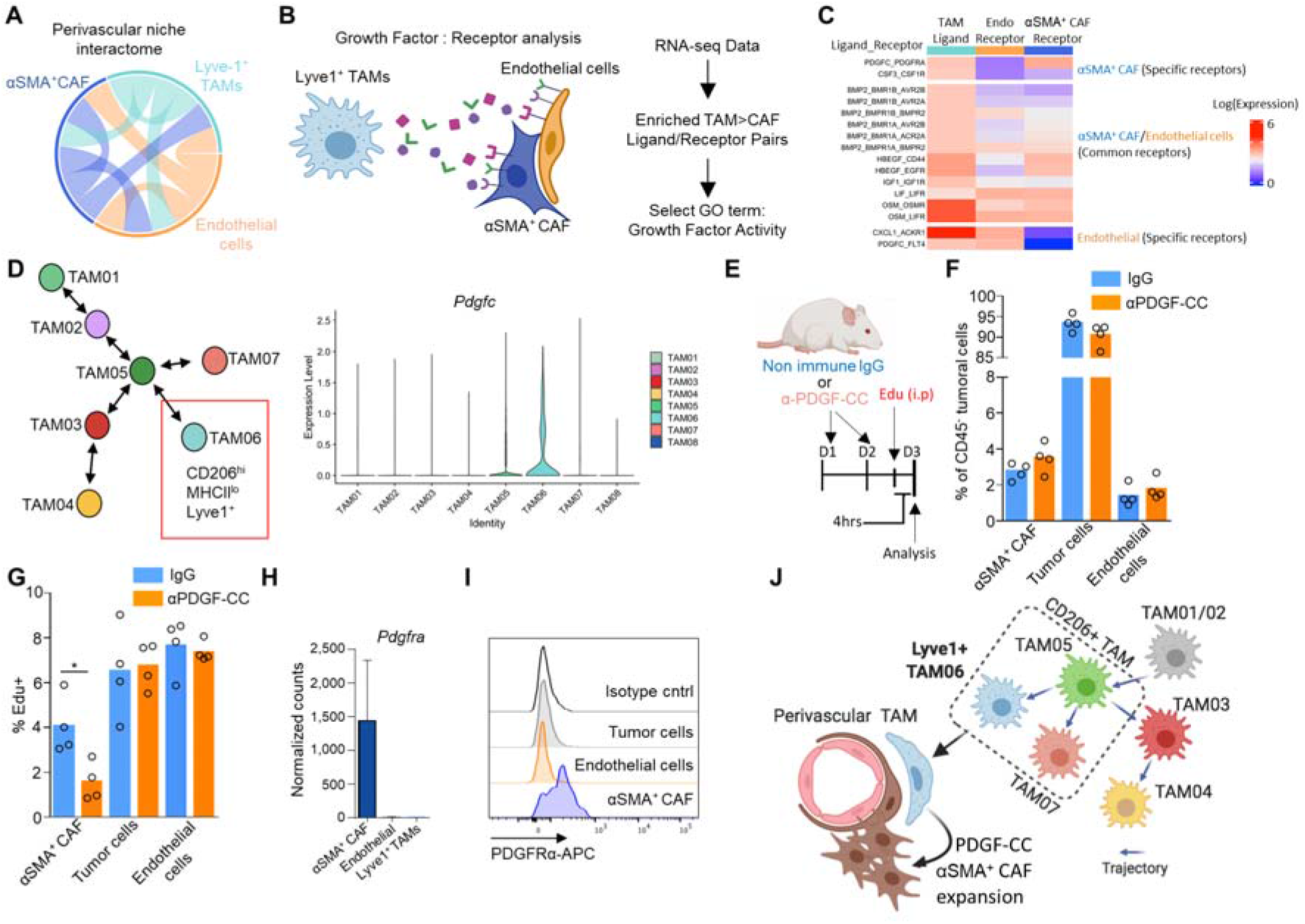
Lyve-1^+^ TAMs communicate to αSMA^+^ CAFs in the perivascular niche via a pro-proliferative PDGF-CC:PDGFR-α interaction. (**A**) Circos plot showing predicted crosstalk of perivascular ligand-receptor interactions as identified by *CellPhoneDB* from the respective RNA-seq datasets. Outer sectors and links between sectors are weighted according to the total number of annotated ligand-receptor interactions between each respective cell type. (**B**) Schematic representing the method of cell type ligand-receptor interactome generation. (**C**) Heatmap showing the Lyve-1^+^ TAM (TAM06) and αSMA^+^ CAF population-specific secretome generated using data from (**A**) and the method outlined in (**B**) diagram displaying the ligand:receptor pairs between Lyve-1^+^ TAMs (TAM06) and αSMA^+^ CAFs and endothelial cells. The analysis highlighted a unique PDGF-CC:PDGFRα interaction specific to Lyve-1^+^ TAMs (TAM06) and αSMA^+^ CAFs. (**D**) Schematic map of each TAM cluster’s location along the respective trajectories marking the Lyve-1^+^ TAM population (left) and violin plots of *Pdgfc* expression associated with TAM clusters (right). (**E-G**) Schematic for experimental approach and dosing strategy to acutely inhibit PDGF-CC signalling using an anti-PDGF-CC neutralizing antibody (**E**). Abundance of indicated cell populations (**F**). Proportion of EdU^+^ cells within each CD45^-^ cell subset, (cohorts of n=4 mice) (**G**). (**H**) Bar plot depicting normalized gene expression values for *Pdgfra* in the bulk RNA-sequenced populations (left) across n=5 mice. (**I**) Representative histograms of surface PDGFRα staining on the indicated cells against isotype antibody staining of gated using flow cytometry analysis from enzyme-dispersed *MMTV-PyMT* tumors. (**J**) Schematic overview of the Lyve-1^+^ TAM supporting niche to support perivascular αSMA^+^ CAF expansion through its close proximity and high expression of PDGF-CC. Images in panel (B and J) was created using *BioRender* software. Bar charts represent mean and the dots show individual data points from individual tumors and mice, error bars represent s.d. * *P*<0.05.

PDGFRs form either homo- or hetero-dimers between the α and β receptor subunit (αα, αβ and ββ) and a homodimer of PDGF-C (PDGF-CC) selectively signals through PDGFRαα and PDGFRαβ dimers (*51, 52*) which has been demonstrated to be a mitogenic and migratory factor for human dermal myofibroblasts (*53, 54*). To assess whether PDGF-CC may play a role in directing the proliferation of the αSMA^+^ CAF population within the perivascular niche we administered neutralizing antibodies to PDGF-CC, within an acute treatment regimen, in tumor bearing *MMTV-PyMT* mice (Fig. 5E). Neutralization of PDGF-CC did not affect the abundance of the cell populations at the acute timepoint (Fig. 5F) but did diminish Edu^+^ incorporation of the αSMA^+^ CAFs, but not in the tumor cells or CD45^-^ CD31^+^ endothelial cells within the vascular niche (Fig. 5G). This highlighted that the expansion of perivascular αSMA^+^ CAFs was PDGF-CC dependent and could account for the role of Lyve-1^+^ pvTAMs in orchestrating expansion of the population during tumor progression. As a population of perivascular fibroblasts have been implicated in recruiting macrophages to the perivascular niche (*13*), these observations in the current study highlight a potential reciprocal interactions between TAMs and mesenchymal populations in perivascular niche formation. PDGF-CC is a prognostic factor for poor survival in breast cancer (*55*) and has been demonstrated to be important to angiogenesis (*56, 57*). Within the perivascular niche the αSMA^+^ CAFs selectively expressed PDGFRα (Fig. 5H,I), alongside PDGFRβ (fig. S4E), and as such, were the only cell to be capable of responding to PDGF-CC. Tumors grow slower in *MMTV-PyMT Pdfgc*^-/-^ mice and display increased necrotic areas and evidence of hemorrhage (*55*). In accordance with our observations in murine models, using The Cancer Genome Atlas (TCGA) we observed an enrichment for a αSMA^+^ CAF signature (using genes identified in the murine population) above that of healthy tissue in human breast cancer (fig. S5C) and interestingly the αSMA^+^ CAF signature also positively correlated with *PDGFC* expression within the tumor (fig. S5D). This study raises an interesting parallel to the observations by Shook et al., that macrophages expressing PDGF-CC support the expansion of αSMA^+^ myofibroblast populations in the wound healing response (*54*), a stromal response which share many similarities to that of cancer (*2, 58*).

This study characterizes a biologically important subset of TAMs selectively expressing Lyve-1. We demonstrate that the Lyve-1^+^ pvTAM subset, which only accounts for 1.4±0.4% of live tumoral cells, is pivotal to tumor growth. We define a new role for pvTAMs in directing the expansion of a perivascular pericyte-like mesenchymal population to form a pro-angiogenic niche that is facilitated by a selective PDGFRα:PDGF-CC crosstalk (Fig. 5J). This study highlights the inter-reliance of stromal populations and the importance of the immune system in orchestrating non-immune stromal cell reactions in cancer which provides therapeutic opportunities for unraveling the complexity of the stromal support network and niches which underpin tumor progression.

## Acknowledgements

The authors thank Dr Yasmin Haque (KCL) and the NIHR BRC flow cytometry platform at Guy’s and St Thomas’ Biomedical Research Centre for cell sorting and flow cytometry assistance, the Nikon Imaging Centre (KCL) for use of their facilities and assistance with confocal microscopy analyses, Dr Alka Saxena (KCL) for support and useful expert discussion regarding the RNA sequencing analyses, Dr Umar Niazi (KCL) for bio-informatic support and Drs Alan Ramsay and Aleksandar Ivetic (KCL) for helpful discussions. We thank Y. Miwa (Tsukuba University), O. Kanagawa (RCAI, RIKEN) and M. Tomura (Osaka Ohtani University) for the Kaede mice. This work was funded by a grant from the European Research Council (335326). P.G. was supported by a grant from the Wellcome Trust (101529/Z/13/2). J.W.O. and J.E.A. are supported by the UK Medical Research Council (MR/N013700/1) and are KCL members of the MRC Doctoral Training Partnership in Biomedical Sciences. PyMT x Kaede studies were supported by a Cancer Immunology Project Award (C54019/A27535) from Cancer Research UK awarded to D.R.W. The research was supported by the Cancer Research UK King’s Health Partners Centre and Experimental Cancer Medicine Centre at King’s College London, and the National Institute for Health Research (NIHR) Biomedical Research Centre based at Guy’s and St Thomas’ NHS Foundation Trust and King’s College London. The views expressed are those of the authors and not necessarily those of the NHS, the NIHR or the Department of Health.

## Author contributions

J.W.O., J.N.A. conceived the project, designed the approach, performed experiments, interpreted the data wrote the manuscript. J.E.A., I.D., E.J.H., I.B., J.C., T.M., P.G., R.N. designed the approach, performed experiments, and interpreted the data. S.E.P., T.N., F.D., S.K., D.R.W., T.L. designed experiments, interpreted the data and provided key expertise.

## Materials and Methods

### Mice

*MMTV-PyMT* (*PyMT*) mice used in this study were on an FVB/N background. Balb/c and C57Bl/6 wild type mice were obtained from Charles River. Female C57Bl/6 homozygous *Kaede* mice (*45*) were crossed with male *MMTV-PyMT* (FVB background) mice and the F1 offspring used experimentally Cohort sizes were informed by prior studies (*2, 8*). All mice used for experiments were female and randomly assigned to treatment groups. Mice were approximately 21-26 g when tumors became palpable. Experiments were performed in at least duplicate and for spontaneous *MMTV-PyMT* tumor studies individual mice were collected on separate days and all data points are presented.

### Tumor studies

Murine 4T1 mammary adenocarcinoma, Lewis lung carcinoma (LL2) and B16-F10 melanoma cells were obtained from ATCC. 2.5 x 10^5^ cells in 100μl RPMI and were injected by subcutaneous (s.c.) injection into the mammary fat pad of syngeneic Balb/c (4T1) or C57Bl/6 (B16-F10 and LL2) female mice that were six to eight weeks of age. In studies using *MMTV-PyMT* mice tumors arose spontaneously. When tumors became palpable, volumes were measured every 2 days using digital caliper measurements of the long (L) and short (S) dimensions of the tumor. Tumor volume was established using the following equation: Volume= (S^2^xL)/2. *PyMT/Kaede* mice were photo-labeled under anesthesia, individual tumors mice were exposed to a violet light (405nm wavelength) through the skin for nine 20 second exposure cycles with a short 5 second break interval between each cycle. Black cardboard was used to shield the rest of the mouse throughout the photoconversion procedure. Mice for 0 h time points were culled immediately after photoconversion. This photoconversion approach was adapted from that used to label peripheral lymph nodes (*59*). Tumor tissue for flow cytometry analyses were enzyme-digested to release single cells as previously described (*41*). In brief, tissues were minced using scalpels, and then single cells were liberated by incubation for 60 mins at 37°C with 1 mg/ml Collagenase I from *Clostridium Histolyticum* (Sigma-Aldrich) and 0.1 mg/ml Deoxyribonuclease I (AppliChem) in RPMI (Gibco). Released cells were then passed through a 70 μm cell strainer prior to staining for flow cytometry analyses. Viable cells were numerated using a hemocytometer with trypan blue (Sigma-Aldrich) exclusion. For drug treatments, drugs were freshly prepared on the day of injection and administered by intraperitoneal (i.p.) injection using a 26 G needle. For EdU experiments mice were injected i.p. with 50 mg/kg EdU dissolved in Dulbecco’s phosphate buffered saline (PBS) and sacrificed 4 hours post-injection. To liposome deplete Pv macrophages, *MMTV-PyMT* mice were injected i.p. with 150 μl of either clodronate- or PBS-filled liposomes (Anionic Clophosome, FormuMax) on the indicated days. To label PvTAM, *MMTV-PyMT* mice were injected i.p. with 150 μl of Dil fluorescent tracing liposomes (Anionic Clophosome, Formumax). To neutralize PDGF-CC, mice were injected i.p. with 100 μg of a goat anti-PDGF-C neutralizing antibody (AF1447, Bio-techne) solubilized in PBS on day −2 and −1 prior to analysis.

### Murine tissue staining

Mouse mammary tumors were fixed overnight (O.N.) in 4% paraformaldehyde, followed by O.N. dehydration in 30% sucrose prior to embedding in OCT and snap freezing in liquid nitrogen. Frozen sections from these tumors were fixed in 4% paraformaldehyde in PBS for 10 mins at RT and were washed in Tris Buffered Saline (100mM Tris, 140mM NaCl), 0.05%, Tween 20, pH7.4 (TBST) and blocked with TBST, 10% donkey serum (Sigma-Aldrich), 0.2% Triton X-100. Immunofluorescence was performed as previously described (*2*). The following antibodies and dilutions were used: F4/80 1:100 (C1:A3-1, Bio-RAD), αSMA 1:100 (AS-29553, Anaspec), CD31 1:100 (MEC13.1, Biolegend), CD31 1:100 (ab28364 Abcam), mKi67 1:100 (AF649 R&D), CD34 1:100 (RAM34, Invitrogen), desmin 1:100 (PA5-19063, Invitrogen). Primary antibodies were detected using antigen specific Donkey IgG, used at 1:200: AlexaFluor^®^ 488 anti-rabbit IgG, AlexaFluor^®^ 488 anti-rat IgG, AlexaFluor^®^ 568 antirabbit IgG, AlexaFluor^®^ 568 anti-goat IgG, AlexaFluor^®^ 647 anti-rabbit IgG (Thermo Fisher Scientific). NL657 anti-rat goat IgG (R&D) and Cy3 anti-sheep donkey IgG (Jackson Immuno) were also used. Viable blood vessels were visualized in mice through i.v. injection of FITC-conjugated dextran (MW20,000, Thermo Fisher Scientific) 20 min prior to sacrifice. Nuclei were stained using 1.25 μg/ml 4’,6-diamidino-2-phenylindole, dihydrochloride (DAPI) (Thermo Fisher Scientific). Images were acquired using a Nikon Eclipse Ti-E Inverted spinning disk confocal with associated NIS Elements software. Quantitative data was acquired from the images using NIS Elements software.

### Human tissue staining

FFPE human breast adenocarcinoma tissue sections of 4 μm were incubated at 60°C for 1 h, before being deparaffinized with Tissue-Tek^®^ DRS^™^2000, Sakura. Heat-induced antigen retrieval was performed using a pressure cooker (MenaPath Access Retrieval Unit, PASCAL). The slides were immersed in modified citrate buffer pH 6 and gradually heated to 125°C. Excess of antigen retrieval buffer was washed firstly with distilled water followed by PBS, before incubation of the slides in blocking buffer containing 0.5% Triton and 5% donkey serum (Sigma) for 30 mins at room temperature. The sections were then probed with anti-CD68 1:100 (KP1, Invitrogen), anti-αSMA 1:200 (1A4, Sigma-Aldrich) and anti-CD31 1:100 (EP3095, Abcam) diluted in blocking buffer overnight at 4°C. After further washing, sections were stained for 2 h with donkey IgG antibodies purchased from Jackson Immunoresearch and used at 1:600; AlexaFluor^®^ 647 anti-mouse IgG and AlexaFluor^®^ 488 anti-rabbit IgG. After washing in PBS, the sections were incubated with anti-αSMA conjugated with CY3 probe 1:200 (1A4, Sigma-Aldrich). Counterstaining was performed with 1:2000 DAPI (Cell Signalling Technology) for 5 mins, followed by a wash step using PBS. Mounting medium (Fluorsave, Millipore) was applied to the slides. Images were acquired using a Nikon Eclipse Ti-E Inverted spinning disk confocal with associated NIS Elements software.

### Flow cytometry

Flow cytometry was performed as previously described (*6*). The following antibodies against the indicated antigen were purchased from Thermo Fisher Scientific and were used at 1 μg/ml unless stated otherwise: CD3ε APC and PE (145-2C11), CD4 FITC (RM4-5), CD8β eFluor®450 (H35-17.2), CD11b APC-eFluor^®^780 (M1/70), CD11b BV510 (M1/70), CD11c APC (N418), CD16/32 (2.4G2; Tonbo Biosciences), CD19 APC (6D5; Biolegend^®^), CD29 APC (eBioHMb1-1), CD31 eFluor^®^ 450 and PE (390), CD34 FITC and APC (RAM34), CD45 APC-eFluor^®^ 780, FITC and PerCP-Cy5.5 (30-F11), CD90.2 eFluor^®^ 450 (53-2.1), CD90.1 eFluor^®^ 450 (HIS51), CD90.1 BV510 (OX-7), CD103 PE (2E7), CD206 APC (FAB2535A; Bio-Techne), F4/80 PE (BM8; Biolegend^®^), F4/80 BV421 (BM8; Biolegend^®^), FAP (10 μg/ml, AF3715, Bio-Techne), Ly6C PE and eFluor^®^ 450 (HK1.4), Ly6G FITC (1A8; Biolegend^®^), Lyve-1 AlexaFluor^®^ 488 (ALY7), MHCII PE and FITC (M5/114.15.2), NK1.1 APC (PK136), PDGFRα PerCP-Cy5.5 (APA5), Ly6A/E AlexaFluor^®^ 700 (D7). Where stated, the following corresponding isotype control antibodies at equivalent concentrations to that of the test stain were used: Armenian Hamster IgG APC (eBio299Arm), goat IgG APC and PE (Bio-techne), rat IgG2a APC, PE and FITC (eBR2a) and rat IgG2b APC and eFluor^®^ 450 (eB149/10H5). Intracellular stains were performed as previously described (*6*). Dead cells and red blood cells were excluded using 1 μg/ml 7-amino actinomycin D (7AAD; Sigma-Aldrich), Fixable Viability Dye eFluor^®^ 780 (Invitrogen) or DAPI alongside anti-Ter-119 PerCP-Cy5.5 or APC-eFluor^®^ 780 (Ter-119; Invitrogen). The FAP primary antibody was detected with a secondary biotin-conjugated anti-goat/sheep mouse IgG and 1:1000 Streptavidin PE-Cy7 (Thermo Fisher Scientific). EdU was detected using the Click-IT Plus Flow Cytometry Assay with AlexaFluor^®^ 488 (Thermo Fisher Scientific) in accordance with the manufacturers’ specifications. Briefly, cells were stained with cell surface antibodies and then fixed and permeabilized and the click chemistry reaction was performed as specified with AF488-conjugated Picolyl Azide to identify EdU incorporated into the genomic DNA. Cells were sorted to acquire pure populations using a FACSAria (BD Biosciences). Data were collected on a BD FACS Canto II (BD Biosciences) or a BD LSR Fortessa (BD biosciences). Data was analyzed using FlowJo software (BD biosciences). Unsupervised clustering of flow cytometry data was performed using the ImmunoCluster package (*60*). Briefly, the single-cell data was asinh transformed with cofactor of 150 and clustering was performed with and ensemble method using FlowSOM (*42*) and ConsensusClusterPlus (*61*) to k=8 clusters, based on the elbow criterion, which were manually merged based on expression profiles into biologically meaningful populations as previously outlined (*62*). Dimensionality reduction for visualization purposes was performed with UMAP (*22*).

### Quantitative real time quantitative PCR

mRNA was extracted from FACS-sorted cell populations using the Trizol method and converted to cDNA/amplified using the CellAmp^™^ Whole Transcriptome Amplification Kit (Real Time), Ver. 2 kit (Takara) according to the manufacturer’s protocol. mRNA of interest was measured using the SuperScriptTM III PlatinumTM One-Step qRT-PCR Kit (Thermo Fisher Scientific) according to the manufacturer’s protocol with the primers/probes *Actb Mm02619580_g1* and *Pdgfc* Mm00480295_m1 (Thermo Fisher Scientific). Expression is represented relative to the housekeeping gene *Actb*. Gene expression was measured using an ABI 7900HT Fast Real Time PCR instrument (Thermo Fisher Scientific).

### Single-cell RNA-sequencing

TAMs (CD45^+^Ly6G^-^CD11b^+^F4/80^hi^) were sorted from enzyme-digested *MMTV-PyMT* tumors and a total of 10,502 TAMs were sequenced from three *MMTV-PyMT* tumors and run through the 10x Genomics Chromium platform. An average of 43k reads per cell, a median of 2,400 genes and median UMI count of 9,491 per cells was obtained. Single-cell suspensions were prepared as outlined in the 10x Genomics Single Cell 3’ V3 Reagent kit user guide (10x Genomics). Briefly, samples were washed with PBS (Gibco) with 0.04% bovine serum albumin (Sigma-Aldrich) and resuspended in 1□ml PBS, 0.04% BSA. Sample viability was assessed using trypan blue (Thermo Fisher Scientific) exclusion and an EVE automated cell counter (Alphametrix) in duplicate, in order to determine the appropriate volume for each sample to load into the Chromium instrument. The sorted TAMs were loaded onto a Chromium Instrument (10x Genomics) to generate single-cell barcoded droplets according to the manufacturers’ protocol using the 10x Genomics Single Cell 3’ V3 chemistry. cDNA libraries were prepared as outlined by the Single Cell 3′ Reagent kit v3 user guide and each of the three resulting libraries were sequenced on one lane each of a HiSeq 2500 (Illumina) in rapid mode.

### Single-cell RNA-sequencing data processing and analysis

The raw sequenced data was processed with the Cell Ranger analysis pipeline version 3.0.2 by 10x Genomics (http://10xgenomics.com/). Briefly, sequencing reads were aligned to the mouse transcriptome mm10 using the STAR aligner (*63*). Subsequently, cell barcodes and unique molecular identifiers underwent Cell Ranger filtering and correction. Reads associated with the retained cell barcodes were quantified and used to build a transcript count tables for each sample. Downstream analysis was performed using the Seurat v3 R package (*64*). Before analysis, we first performed quality control filtering with the following parameters: cells were discarded on the following criteria: where fewer than 800 unique genes detected, reads composed greater than 12% mitochondrial-associated gene transcripts and cells whose number of reads detected per cell was greater than 65k for sample 1 and 2, 60k for sample 3. All genes that were not detected in at least ten single cells were excluded. Based on these criteria the final dataset contained 9,615 TAMs with 25,142 detected genes. The data was first normalized using the LogNormalize function and a scale factor of 10,000. The 2,000 genes with highest variance were selected with the FindVariableGenes function. In order to minimize the effect of cell cycle associated genes in the dimensionality reduction and clustering, cell cycle associated genes defined by the GO term ‘Cell Cycle’ were removed from the variable gene dataset resulting in 1,765 variable genes. Principal component (PC) analysis was used on the highly variable genes to reduce the dimensionality of the feature space and 35 significant PCs were selected for downstream analysis. To reduce biases caused by technical variation, sequencing depth and capture efficiency, the three sequencing samples were integrated using the Seurat integration method (*64*) as specified. Clusters were identified by a graph based SNN clustering approach within Seurat using the resolution parameters 0-1 in steps of 0.1, followed by analysis using the Clustree R package (*65*). Finally, we used resolution parameter of 0.4 to define 10 clusters. Differentially expressed genes were identified using the FindAllMarkers function where the genes must be detected in a minimum of 25% of cells and have a logFC threshold of 0.25. After identifying marker genes, we excluded two clusters which contained suspected contaminating epithelial cells (enriched in *Epcam, Krt18, Krt8*) and dying low-quality cells (enriched in mitochondrial genes and ribosomal subunit genes). Ultimately, we identified 8 relevant clusters. We used the Slingshot R package (*23*) to investigate inferred developmental trajectories in our TAM population. Briefly, dimensionality reduction was performed using diffusion maps with the Destiny R package (*54*) using the significant PCA principal components used for clustering. A lineage trajectory was mapped into the diffusion space using the first 15 diffusion components (DCs) by Slingshot and each cell was assigned a pseudotime value based on its predicted position along the predicted trajectories. We selected the cluster TAM01 as the base state for the trajectory because it had the lowest M1/M2 activation-associated gene score amongst the terminal trajectory branch clusters, no discriminating upregulated GO pathways and the fewest differentially expressed genes and represented the most naïve TAM transcriptomic base state. To detect non-linear patterns in gene expression over pseudotime trajectory, we used the top variable gene set and regressed each gene on the pseudotime variable we generated, using a general additive model (GAM) with the GAM R package (*37*). Heatmaps were generated with the ComplexHeatmap package (*46*).

### Bulk RNA-sequencing

Cells were sorted directly into RLT plus buffer (Qiagen) supplemented with 2-β-mercaptoethanol (BME) (Gibco) and lysates were immediately stored at −80°C until used. RNA was extracted with the RNeasy Plus Micro kit (Qiagen) as per the manufacturers’ protocol, in addition to on-column DNase digestions specified by the manufacturer (Qiagen). cDNA was generated and amplified using the SMARTseq v4 Ultra Low Input RNA Kit (Clontech) on the contactless Labcyte liquid handling system (Beckman Coulter Life Sciences). Two hundred ng of amplified cDNA was used from each sample where possible to generate libraries using the Ovation Ultralow Library System V2 kit (NuGEN). In brief, cDNA was fragmented through sonication on Covaris E220 (Covaris Inc.), repaired, and polished followed by ligation of indexed adapters. Adapter-ligated cDNA was pooled before final amplification to add flow cell primers. Libraries were sequenced on HiSeq 2500 (Illumina) for 100 paired-end cycles in rapid mode.

### Bulk RNA-sequencing data processing and analysis

Pre-alignment QC for each sample, independently for forward and reverse reads, was performed using the standalone tool FastQC. Reads were trimmed using Trimmomatic (*66*) and aligned to the reference genome (mm10) using HISAT2 (*67*). PCR duplicates were removed using SAMtools (*68*). Counts were generated using the GenomicAlignment (*69*) package using mm10. Prior to performing differential gene expression analysis, genes with very low expression were discarded. Differential expression analysis was performed with DESeq2 (*70*) package in R. The test statistics’ p-values were adjusted for multiple testing using the procedure of Benjamini and Hochberg. Genes with adjusted p-values lower than 0.05 and absolute log2 fold change greater than 1 were considered significant.

### Gene ontology pathway enrichment analysis

Enriched pathways were identified based on differentially expressed genes using gProfiler(*71*) (http://www.biit.cs.ut.ee/gprofiler/). We used pathways gene sets from the ‘biological processes’ (GO:BP) of Gene Ontology (http://www.geneontology.org/). All p-values were adjusted for multiple testing using the procedure of Benjamini and Hochberg.

### Ligand:receptor mapping analysis

Ligand:receptor mapping was performed with the online implementation of the CellPhoneDB v1.0 tool (https://www.cellphonedb.org/) (*72*) run without the statistical method. Cell type ligand:receptor interactome was generated with bulk RNA-seq data as input, selecting genes with expression of 16 normalized counts or greater as input. The resulting interaction list was filtered by selecting non-integrin mediated interactions and TAM ligands that were enriched in the TAM06 scRNA-seq population in the ligand:receptor pairs, finally selecting for ligands present in the GO term ‘growth factor activity’ that were investigated further as potential candidates.

### Computational analysis of cancer patient data

RSEM normalized expression datasets from the Cancer Genome Atlas (TCGA) were downloaded from the Broad Institute Firehose resource (https://gdac.broadinstitute.org/) and analyzed using custom R scripts. The CAF1 gene expression signature was generated by taking the mean normalized log2-transformed expression value of the component signature genes. The CAF1 gene signature genes were selected from the top 25 differentially expressed CAF1 genes by Log Fold change as the maximum set for which a significant positive correlation was observed between all genes and *ACTA2* (αSMA). The final gene set was as follows: *ACTA2, MMP13, LRRC15, COL10A1, SPON1, COL1A1*.

### Statistics

Normality and homogeneity of variance were determined using a Shapiro-Wilk normality test and an F-test respectively. Statistical significance was then determined using a two-sided unpaired Students *t* test for parametric, or Mann-Whitney test for nonparametric data using GraphPad Prism 8 software. A Welch’s correction was applied when comparing groups with unequal variances. Statistical analysis of tumor growth curves was performed using the “compareGrowthCurves” function of the statmod software package (*73*). No outliers were excluded from any data presented.

### Study approval

All experiments involving animals were approved by the Animal and Welfare and Ethical Review Boards of King’s College London and the University of Birmingham, and the Home Office UK. Human breast adenocarcinoma tissue was obtained with informed consent under ethical approval from the King’s Health Partners Cancer Biobank (REC reference 12/EE/0493).

### Data availability

The transcriptomic datasets that support the findings of this study will be made available through the Gene Expression Omnibus. The authors declare that all other data supporting the findings of this study are available within the paper and its supplementary information files.

