## Supplemental_figures_S1-S5 for "Macrophages orchestrate the expansion of a pro-angiogenic perivascular niche during cancer progression"

### Supplementary Figures

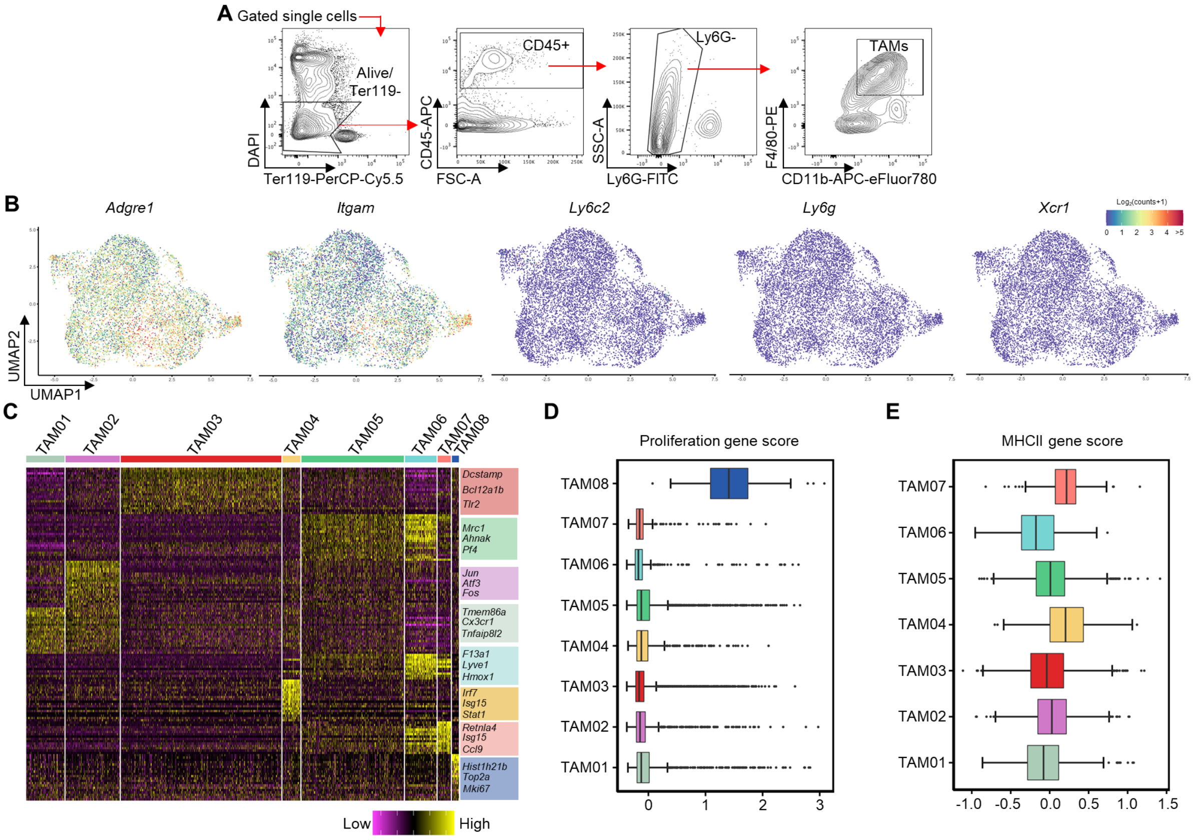


**Figure S1. scRNA-seq of TAMs identifies distinct polarization states within the tumor microenvironment.** (**A**) Representative gating strategy for live (7AAD$^{-}$) TAMs for sorting using the indicated surface markers from enzyme-dispersed *MMTV-PYMT* tumors used in scRNA-seq sample preparation analyzed in Fig. 1 and 2. (**B**) UMAP plots of the 9,039 TAMs sequenced using the 10X Genomics' Chromium platform across n=3 tumors displaying the expression of marker genes used to isolate TAM single cells in (**A**). (**C**) Heatmap showing top differentially expressed genes within each TAM cluster shown in Fig.1b, selected genes for each cluster are highlighted to the right of the heatmap. (**D,E**) Gene scores across all TAM clusters for proliferation (**D**) and MHCII associated genes (**E**). Box and whisker plots, the boxes show median and upper and lower quartiles and whiskers shows the largest value no more than 1.5*IQR of the respective upper and lower hinges, outliers beyond the end of the whisker are plotted as individual dots.

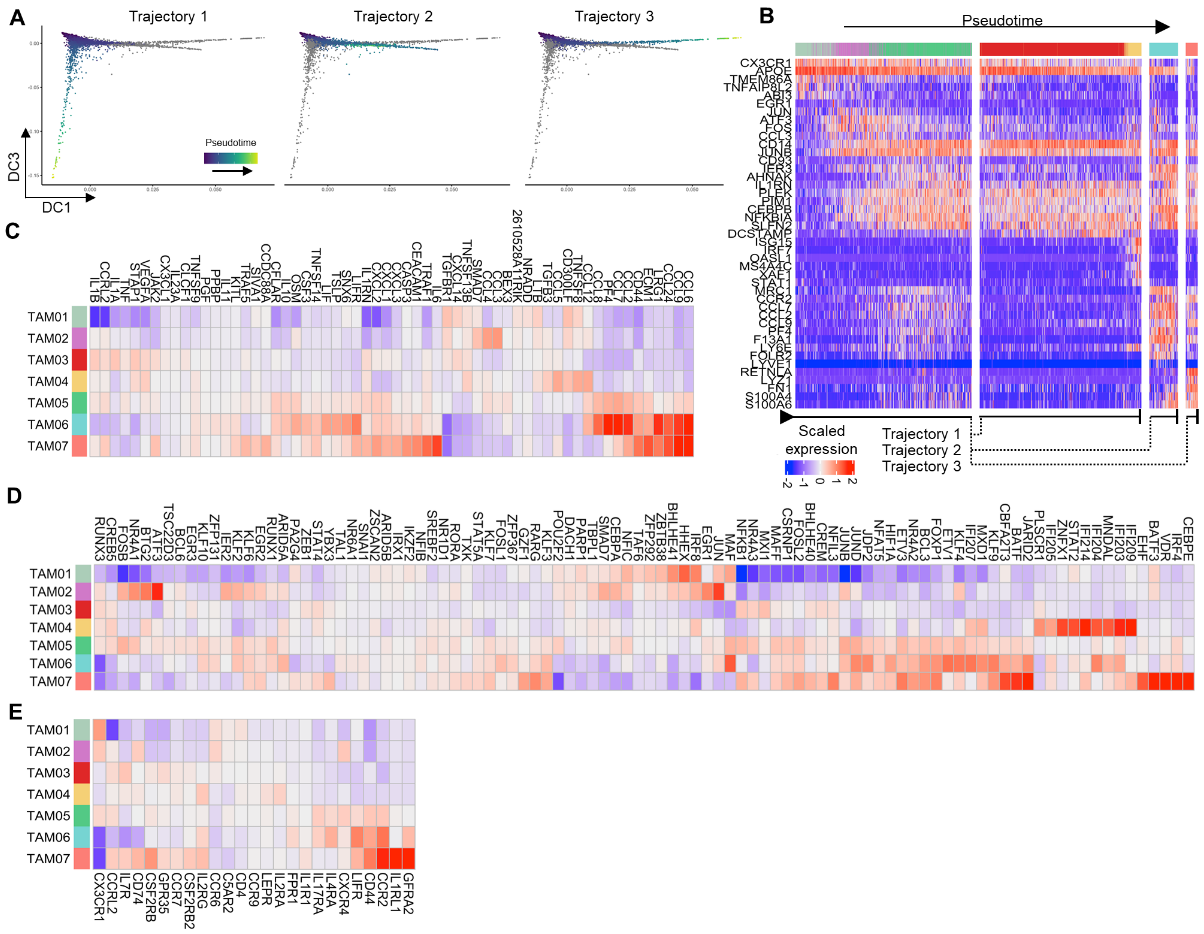

**Figure S2. Trajectory analysis of scRNA-seq of TAMs reveals polarization-specific transcription factors and cytokine signatures associated with transcriptomic state**. (**A**) Diffusion component (DC) plot showing single cells colored by pseudotime value for each distinct trajectory predicted by *Slingshot* trajectory analysis. Cells that are not associated with a given trajectory are greyed out. (**B**) Heatmap of selected genes varying significantly across trajectory. Single cells (columns) are arranged left to right by pseudotime value within their labeled trajectories, branching points are indicated beneath the heatmap. (**C-E**) Heatmaps representing chemokine, cytokine and complement factor genes (**C**), transcription factor genes (**D**) and additional chemokine, cytokine and complement receptor genes (**E**) that vary significantly across the predicted *Slingshot* trajectories.

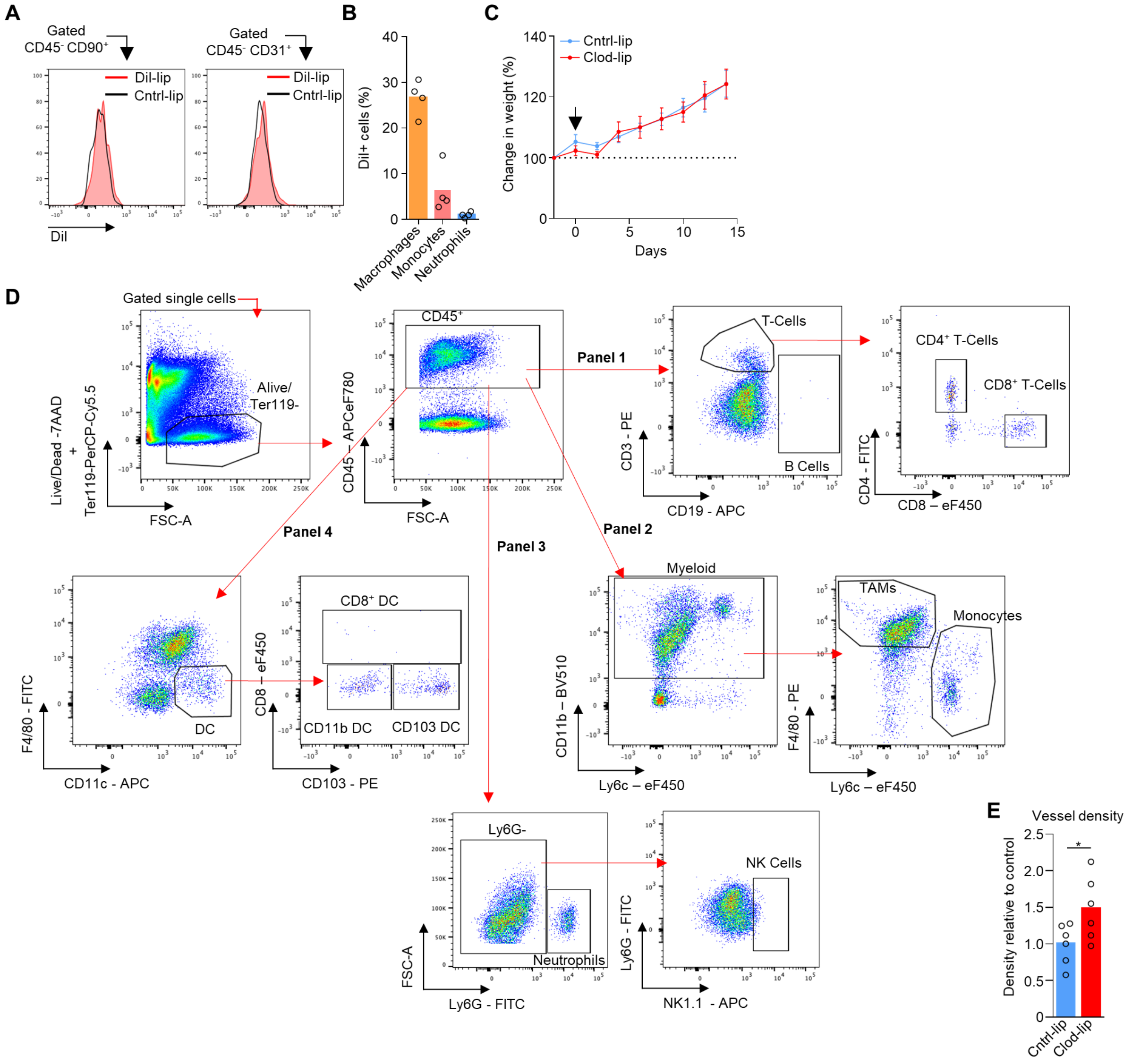


**Figure S3. Liposomes are specifically taken up by a PvTAM subset of TAM.** (**A-B**) Representative histograms showing Dil-containing liposome uptake (red filled histogram) against PBS-filled liposome treated mice (open black line histogram) in CD90$^{+}$ CAF and CD31$^{+}$ endothelial cells in the CD45$^{-}$ gate of enzyme-dispersed tumors from *MMTV-PyMT* mice treated as shown in Fig. 2J and assessed using flow cytometry, representative of n=3 mice (**A**)**.** Proportion of live (7AAD$^{-}$) gated tumor-resident phagocytic populations with detectable Dil uptake as assessed using flow cytometry (n=4 mice) (**B**). (**C-E**) Tumor bearing *MMTV-PyMT* mice treated with control PBS-filled liposomes (Cntrl-lip) or clodronate-filled liposomes (Clod-lip) using the treatment regimen shown in Fig. 3A, showing the percentage change in mouse weight over the course of the experiment (**C**), representative gating strategy for identifying different tumour-resident leukocyte populations (**D**), and number of distinct CD31$^{+}$ endothelial vessel elements as assessed using immunofluorescence analysis stained frozen tissue sections (a total of n=12 sections, across n=6 mice per condition) (**E**). Bar charts represent mean and the dots show individual data points from individual tumors and mice. * *P*<0.05, ** *P*<0.01.

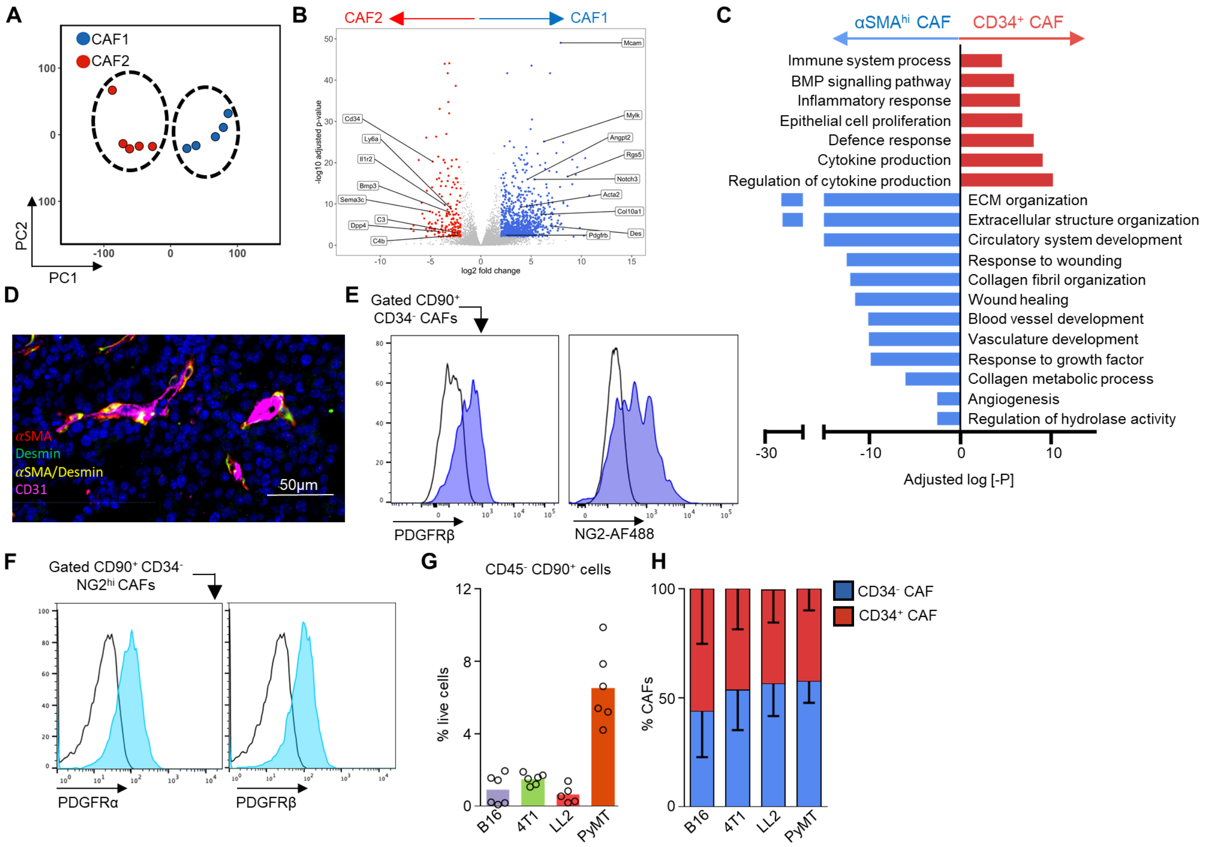


**Figure S4. CD34$^{+/-}$ CAF subsets are transcriptionally distinct and conserved across other murine models of cancer.** (**A-C**) Bulk RNA-seq of the CD34$^{+/-}$CAF subsets, showing PCA plot of the bulk-sequenced CAF populations showing the difference in CAF transcriptome (**A**), differentially expressed genes between CD34$^{+/-}$ CAF populations (**B**) and significantly upregulated GO terms based on differentially expressed genes of the two CAF subsets (**C**), across n=5 tumors and mice. (**D**) Representative confocal image of frozen *MMTV-PyMT* tumor sections showing DAPI (nuclei; blue), and antibody staining against CD31 (magenta), αSMA (red), desmin (green) and αSMA/desmin co-localization (yellow). Representative of multiple sections from n=4 tumors and mice. Scale bar represents 100µm. (**E-F**) Representative histogram of live (7AAD$^-$) CD90$^+$CD34$^-$ CAFs gated using flow cytometry in a representative enzyme-dispersed MMTV-PyMT tumor; histogram shows surface staining for PDGFRβ or NG2 as indicated (blue shaded) against that of the isotype control (open) (**E**), or gated live (7AAD$^-$) CD90$^+$CD34$^-$ NG2$^{hi}$ CAFs showing surface staining for PDGFRα and PDGFRβ as indicated (light blue shaded) against that of the isotype control (open). (**G-H**) Mice were injected with the indicated tumor cells and when tumors reached 1500mm$^3$ they were enzyme-dispersed and analyzed by flow cytometry for the abundance of live (7AAD$^-$) CD45$^-$CD31$^-$CD90$^+$ CAFs (**G**) and the relative proportions of CD34$^+$ and CD34$^-$ CAFs (**H**) across n= 5-6 mice per model. Bar charts represent mean and error bars s.d., dots show individual data points from individual tumors and mice.

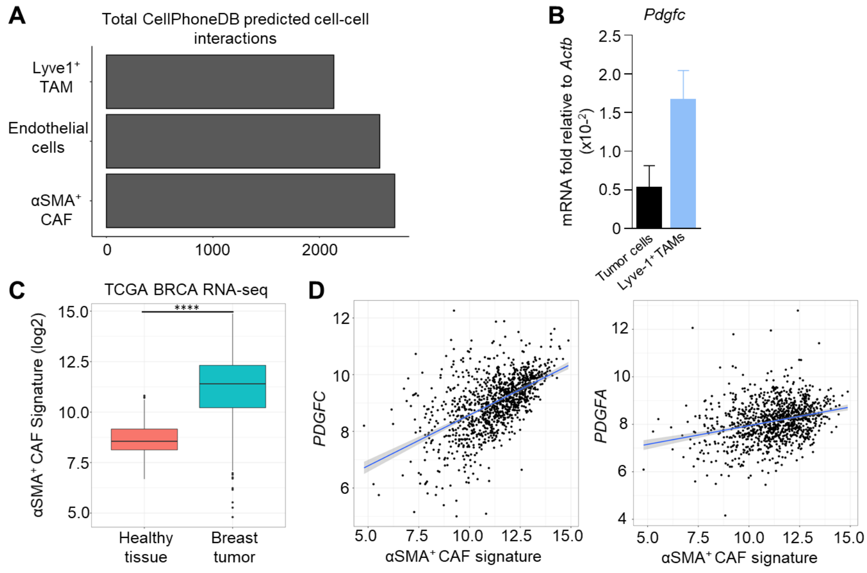


**Figure S5. A αSMA$^+$ CAF signature correlates with *PDGFC* expression in human breast cancer** (**A**) The number of total potential ligand and receptor interactions identified per bulk RNA-sequenced cell type that participate in paracrine and autocrine signaling networks in the perivascular niche. (**B**) *Pdgfc* mRNA expression relative to the housekeeping gene *Actb* in FACS-sorted tumor cells (CD45$^-$CD31$^-$CD90$^-$; n=6) and Lyve-1$^+$ TAMs (n=3). (**C**) Normalized log2 RNA-seq counts for the TCGA-BRCA dataset for the αSMA$^{hi}$ CAF gene signature in non-malignant breast tissue (n=112) and primary breast carcinoma tissue (n=1,093). (**D**) Scatterplot of αSMA$^+$ CAF gene signature score (x axis) and *PDGFC* (y axis) (Pearson's r = 0.547, p < 0.0001) (left) or *PDGFA* (y-axis) (Pearson's r = 0.286 p < 0.0001) (right) from the TCGA-BRCA RNA-seq dataset, n=1,093. Box and whisker plots, the boxes show median and upper and lower quartiles and whiskers shows the largest value no more than 1.5*IQR of the respective upper and lower hinges, outliers beyond the end of the whisker are plotted as individual dots. Bar charts represent mean, error bars represent s.e.m. **** *P*<0.0001.
